# Longitudinal increases in structural connectome segregation and functional connectome integration are associated with better recovery after mild TBI

**DOI:** 10.1101/320515

**Authors:** Amy F. Kuceyeski, Keith W. Jamison, Julia P. Owen, Ashish Raj, Pratik Mukherjee

## Abstract

Traumatic brain injury damages white matter pathways that connect brain regions, disrupting transmission of electrochemical signals and causing cognitive and emotional dysfunction. Connectome-level mechanisms for how the brain compensates for injury have not been fully characterized. Here, we collected serial MRI-based structural and functional connectome metrics and neuropsychological scores in 26 mild traumatic brain injury subjects (29.4±8.0 years, 20 male) at 1 and 6 months post-injury. We quantified the relationship between functional and structural connectomes using network diffusion model propagation time, a measure that can be interpreted as how much of the structural connectome is being utilized for the spread of functional activation, as captured via the functional connectome. Overall cognition showed significant improvement from 1 to 6 months (t_25_=−2.15, p=0.04). None of the structural or functional global connectome metrics were significantly different between 1 and 6 months, or when compared to 34 age- and gender-matched controls (28.6±8.8 years, 25 male). We predicted longitudinal changes in overall cognition from changes in global connectome measures using a partial least squares regression model (cross-validated R^2^ = 0.27). We observe that increased network diffusion model propagation time, increased structural connectome segregation and increased functional connectome integration were related to better cognitive recovery. We interpret these findings as suggesting two connectome-based post-injury recovery mechanisms: one of neuroplasticity that increases functional connectome integration and one of remote white matter degeneration that increases structural connectome segregation. We hypothesize that our inherently multi-modal measure of network diffusion model propagation time captures the interplay between these two mechanisms.

**Abbreviations:** mild traumatic brain injury (mTBI), structural connectome (SC), functional connectome (FC), network diffusion (ND), functional MRI (fMRI), diffusion MRI (dMRI), principal component analysis (PCA), partial least squares regression (PLSR), confidence interval (CI), Attention Network Test (ANT), California Verbal Learning Test II (CVLT-II), Coma Recovery Scale – Revised (CRS-R)

## 1.0 Introduction

Impaired cognitive abilities, particularly attention and memory, are the most common and debilitating cognitive deficits following traumatic brain injury (TBI) [Ashman et al., 2006; Brenner, 2011; Willmott et al., 2009]. More than 5.3 million persons in the US alone are living with TBI-related cognitive dysfunction [Langlois et al., 2006], with an estimated 1.5 million new cases each year, resulting in a total annual medical cost of $77 billion. Recent increases in sports-related and military-related mild TBI (mTBI) have propelled research focus on this disease to the forefront. While both spontaneous and rehabilitation-driven recovery is observed in some individuals after mild TBI [McCrea et al., 2009], between 10-20% of sufferers have persistent cognitive or emotional dysfunction [McAllister et al., 2006; Wood, 2004].

Diffuse axonal injury that occurs in mTBI can result in neurological impairment by damaging the brain’s structural white matter connections, impacting their ability to transmit neuronal signals. Many studies have found relationships between biomarkers of diffuse axonal injury, including diffusion tensor imaging summary statistics such as fractional anisotropy, and cognitive impairment [Kuceyeski et al., 2011; Niogi et al., 2008; Sharp et al., 2014; Yuh et al., 2014]. Diffuse axonal injury can also impact network-level measures of structural connectivity (SC) and functional connectivity (FC). Connectomics, a method that enables network-level analysis of anatomical (measured via diffusion MRI) and physiological (measured via functional MRI, magnetoencephalography or EEG) connections between brain regions, has also been applied in a range of neurological disorders, including mTBI [Chu et al., 2018a; Irimia et al., 2012; Sharp et al., 2014; Spielberg et al., 2015]. Pandit et al., (2013) found reduced overall FC, longer global characteristic path length and reduced global efficiency in mTBI vs. normal controls, while Nakamura et al., (2009) showed lower small world indices in the resting-state FC network. A few publications report improvements in SC related to recovery from severe brain injury [Fernández-Espejo et al., 2011; Sidaros et al., 2008; Voss et al., 2006], although another showed long-term impairment of white matter 5 years post-injury even in those individuals that had recovered [Dinkel et al., 2014]. Correlations between network-level improvements in FC and recovery post-injury are more widely reported [Demertzi et al., 2014; Laureys and Schiff, 2012; Sharp et al., 2011; Soddu et al., 2011; Vanhaudenhuyse et al., 2010].

While recovery from TBI depends on both the pattern of initial or continued damage to the SC network and plasticity of the SC and FC network, few studies analyze the two together [Caeyenberghs et al., 2013; Caeyenberghs et al., 2017]. One study showed TBI patients with more SC injury had less FC in the default mode network [Sharp et al., 2011]. Palacios et al., (2013) found increased FC in frontal areas in chronic TBI patients compared to controls, which was also positively correlated with better cognitive outcomes and negatively correlated with SC measures.

While these studies shed light on the relationship between function, structure and recovery, they are statistical or phenomenological, and do not utilize latest advances in modeling the relationship between functional and structural connectomes. Recent work has focused on implementing mathematical models that formalize the relationship between SC and FC in both normal and pathological populations [Cabral et al., 2011; Chu et al., 2018b; Das et al., 2014; Deco et al., 2012; Fernández Galán, 2008; Honey et al., 2009; Messé et al., 2014; Woolrich and Stephan, 2013]. Some of the main goals in joint structure-function modeling are to increase the accuracy of noisy connectivity measurements, identify function-specific subnetworks [Chu et al., 2018b] or to predict one modality from the other [Honey et al., 2009]. One recent publication with the goal of predicting function from structure used the Network Diffusion (ND) model [Abdelnour et al., 2014; Abdelnour et al., 2018], which assumes functional activation diffuses along white matter connections. This model is linear, has a simple, closed-form solution and only one tuning parameter, making it computationally more tractable and less prone to overfitting than, for example, highdimensional, non-linear neural mass models. The ND model has been applied to predicting patterns of atrophy in dementia [Raj et al., 2012], epilepsy [Abdelnour et al., 2015] and a range of neurological disorders[Cauda et al., 2018]. The ND model’s one tuning parameter, called ND model propagation time, allows quantification of the FC-SC relationship. ND model propagation time can be interpreted as the amount of model time that is needed to “play out” the simulated activation propagation within the SC network to best match the observed FC, i.e., “network depth”. ND model propagation time can be interpreted as a metric of “distance” between SC and FC; the smaller the distance, the more trivially FC can be explained by SC. Our recent cross-sectional study showed that ND model propagation time was the only global connectome metric (including FC and SC metrics) correlated with Coma Recovery Scale – Revised (CRS-R) after severe brain injury [Kuceyeski et al., 2016]. Specifically, increased ND model propagation time was correlated with better level of consciousness as measured by CSR-R, a finding that we also replicated in numerical simulations of injury and recovery. In that work, we interpreted the observed relationship between increased ND model propagation time and better recovery as evidence for global cerebral neuroplasticity in recovery, i.e. reorganized functional connections to compensate for irrevocably damaged structural connections. Here, we test for multi-modal connectomic reorganization mechanisms by examining if longitudinal increases in ND model propagation time, as well as other global functional and structural connectome measures, are associated with better cognitive recovery in mild TBI patients, using tests of attention and memory (see Figure 1). If our multi-modal connectomic measures can predict longitudinal recovery, it will enable a highly parsimonious and potentially clinically relevant biomarker of the mechanism of cognitive recovery post-mTBI. Studies in this population have historically been beset with small effect sizes, diffuse and subtle structural effects and heterogeneous presentations. To our knowledge, this type of model-based exploration of the longitudinal evolution of the connectomes has not before been reported in mTBI.

**Figure 1:**
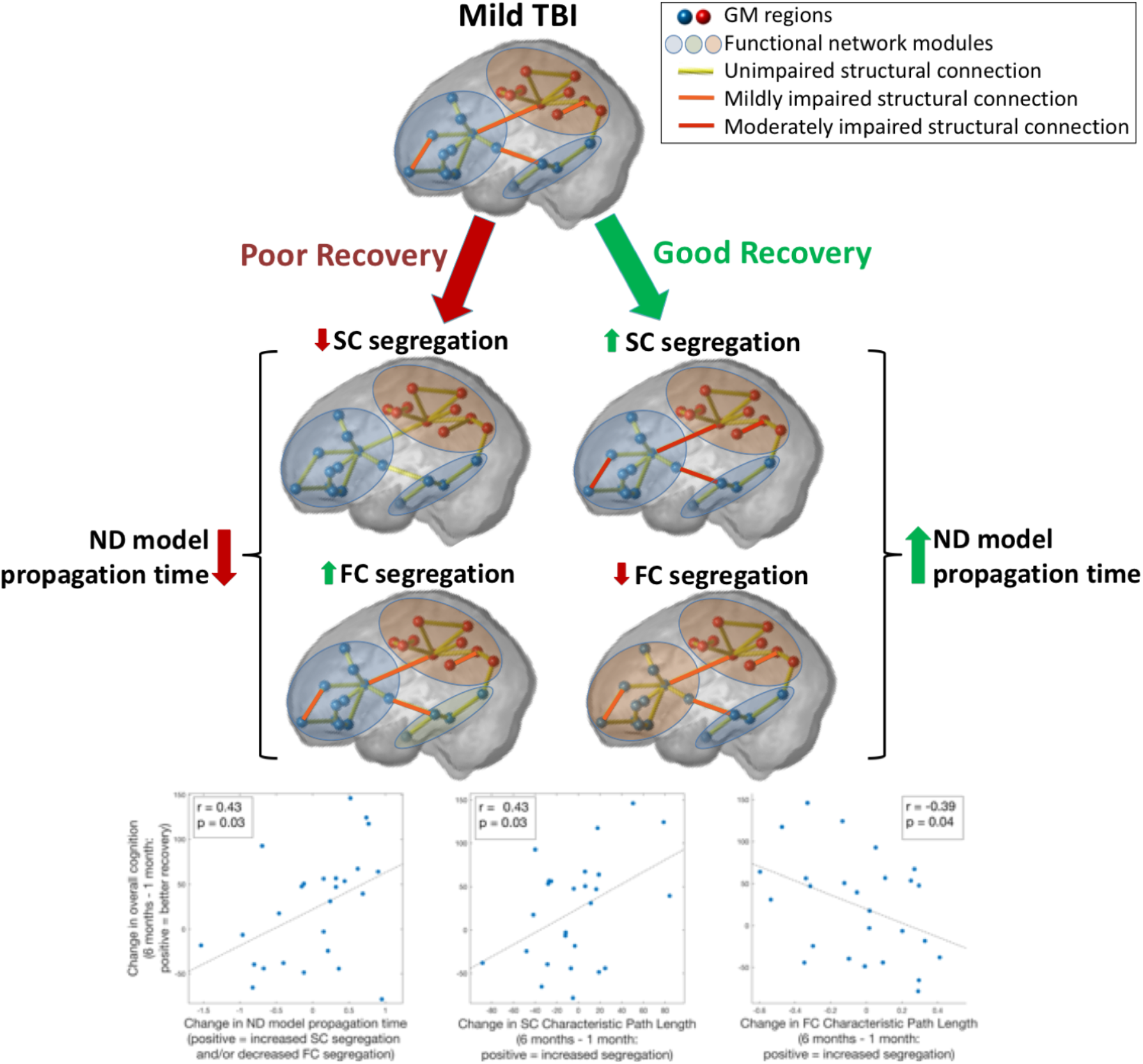
Increased Network Diffusion (ND) model propagation time in overall cognitive recovery after mild TBI. We hypothesize that the interval increase in ND model propagation time is reflecting two processes that track with recovery: increased structural segregation and decreased functional segregation. The bottom three panels show the univariate correlations between change in recovery measures from one to 6 months and 1) change in ND model propagation time, 2) change in structural connectome segregation (characteristic path length) and 3) change in functional connectome segregation (characteristic path length); Pearson correlation and uncorrected p-values are given in the panel inserts.

## 2.0 Materials and Methods

### 2.1 Data

Data from 51 subjects (29.6±8.6 years of age, 35 male) that incurred mild TBI was collected at 1 week, 1 month, 6 months and 12 months post-injury. Our hypothesis is that most recovery occurs between the 1 month and 6 month time points [Losoi et al., 2016], so we chose to focus on the data from those two time points only. A total of 27 subjects had complete datasets (neuropsychological test scores and MRI data) from both 1 and 6 months (29.1±8.1 years of age, 21 male). The conditions for inclusion were blunt, isolated mTBI, defined as Glasgow Coma Scale of 13–15 at injury, loss of consciousness less than 30 minutes and post-traumatic amnesia less than 24 hours. No imaging was used to define mTBI. The conditions for exclusion were pregnancy or other contraindication to MRI, a history of neurological/psychiatric diagnosis, prior seizure, or drug/alcohol abuse. MRIs were collected on a 3T GE Signa EXCITE scanner and included structural scans (FSPGR T1, 1×1×1 mm^3^ voxels), resting-state fMRI (7 min, 3.4×3.4×4.0 mm^3^ voxels, 2 sec sampling rate) and 55-direction high angular resolution diffusion MRI (dMRI: b=1000 sec/mm^2^, 1.8x×1.8×1.8 mm voxels). Neuropsychological testing of attention and learning/memory consisted of 9 subscores within the Attention Network Test (ANT) [Fan et al., 2002], as well as 16 subscores of the California Verbal Learning Test-II (CVLT-II) [Jacobs and Donders, 2007], including measures of short delay/long delay/free/cued recall and total intrusions/repetitions/recognition. The same MRI sequences were acquired in 34 age- and gender-matched normal controls (28.6±8.8 years, 25 male) for comparison.

### 2.2 Image Processing

Gray matter and white matter tissues were classified and the gray matter further parcellated into 86 anatomical regions of interest using the semi-automated FreeSurfer software [Fischl and Dale, 2000]. Cortical and subcortical parcellations were then used in the construction of the SC and FC networks.

#### 2.2.1 Extraction of the functional connectomes

All data were analyzed in Matlab using SPM12 and the CONN functional connectivity toolbox 17f (http://www.nitrc.org/projects/conn) [Whitfield-Gabrieli and Nieto-Castanon, 2012]. Preprocessing of the fMRI data was performed using the CONN toolbox “Direct normalization to MNI-space” pipeline, which includes motion-correction (simultaneous realignment and unwarping), slice-timing correction, and coregistration/normalization to MNI space (3mm voxels). Outlier volumes were removed automatically using the Artifact Detection Tools within the CONN toolbox. The toolbox performs a rigorous regression of head motion (24 total motion covariates: 6 motion parameters plus temporal derivatives and squared terms) and physiological artifacts (10 total CompCor eigenvariates: 5 each from eroded WM and CSF masks [Behzadi et al., 2007]). Notably, this denoising does not regress out global signal, allowing for interpretation of anti-correlations [Chai et al., 2012]. Band-pass filtering (0.008-0.09 Hz) of the residual blood oxygen level-dependent contrast signal was also conducted. Each subject’s cortical and subcortical parcellation from FreeSurfer was coregistered and transformed into MNI space, and these parcels were used to extract average functional time series for each anatomical region of interest. The pairwise FC between two regions was defined as the Pearson correlation coefficient between these time-dependent regional signal averages after removing the first five volumes. Correlation coefficients with a corrected p-value of greater than 0.05 were set to zero. Correction for multiple comparisons was performed for each individual using the linear step-up procedure for false discovery rate correction introduced in [Benjamini and Hochberg, 1995].

#### 2.2.2 Extraction of the structural connectomes

Diffusion MRIs were linearly motion corrected using a modified version of FSL’s eddy_correct and the linear correction applied to the gradient directions. The dMRIs were then corrected for eddy currents using FSL’s eddy_correct. Orientation distribution functions were constructed using FSL’s bedpostx (2 fiber orientations, 1000 sample burn in), gray/white matter masks linearly transformed to dMRI space and streamline tractography performed from each voxel in the gray matter/white matter interface (linear interpolation, Euler tracking, step size = 0.625, threshold for fractional anisotropy > 0.15, curve threshold = 70, curve interval = 2). SC matrices were calculated as the number of streamlines connecting any given pair of regions.

### 2.3 Global Connectome Metrics

Global metrics of average degree, characteristic path length, global and local efficiency, clustering coefficient, modularity, small-world index, transitivity, average mean first passage time [Goñi et al., 2013], mean navigation time (based on both connectivity strength and Euclidean distance) [Seguin et al., 2018] were calculated for the weighted SC and FC networks using the Brain Connectivity Toolbox [Rubinov and Sporns, 2010]. The ratio of between to within-module connection strength was taken to be the ratio of the average strength of edges between nodes in different modules divided by the average strength of edges between nodes in the same module. Before calculation of clustering coefficient, local efficiency and transitivity, entries in the connectomes were rescaled between 0 and 1 by dividing each entry in the matrix by the maximum value. Negative entries in the FC matrices were removed for connectome metric calculations. Each edge in the SC matrices was divided by the sum of the volumes of the two regions, allowing correction for different sized regions that would have proportionally more/fewer number of seeds in the tractography algorithm. It also adjusts the patient SC to account for any damage-related atrophy in the gray matter regions, allowing for better comparison of graph theoretical measures, since the normalized connection strength is a measure of amount of connectivity proportional to the amount of gray matter that remains. Small-world index s was calculated as

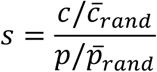

where *c* and *p* are the clustering coefficient and characteristic path length, respectively, of the individual’s network. The variables 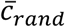 and 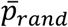 are the means of the clustering coefficient and characteristic path length values, respectively, of 100 different matrices, each obtained by randomly permuting the original connectivity network’s edges 10 times while preserving degree distribution.

### 2.4 Network Diffusion Model Propagation Time

The network diffusion model, detailed in Abdelnour *et al.*, (2014), relates FC to SC by assuming that neuronal activity (functional activation) diffuses within the SC network. In other words, functional activation is modeled as a random walk within the SC network. Therefore, the rate of change of activation at any node *i*, denoted *x_i_*, is related to the difference between the level of activation at that node and its connected neighbors, relative to the sum of outgoing connections of each node (node degree). That is,

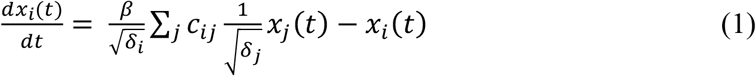

where the coefficients *c_ij_* are the elements of the SC matrix C, *δ_i_* = Σ*_j_c_ij_* is the degree of node *i* and *β* is the rate constant of the exponential decay. This relationship is extended to the entire brain network *x*(*t*)

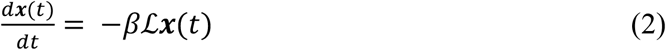

where 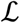 is the well-known network Laplacian. The network Laplacian can have different formulations depending on the normalization factor. We choose, as in Abdelnour *et al.*, (2014) and Kuceyeski *et al.*, (2016), to normalize by node degree, resulting in the Laplacian 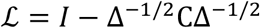, where Δ is the diagonal matrix with entries *δ_i_*. We chose to normalize by node degree in order to control for different sized regions in the gray matter parcellation. Therefore, the matrix C in the calculation of the Laplacian is the SC matrix based on streamline count. For any initial configuration, or activation pattern, ***x_0_***, the solution to Eq (2) is:

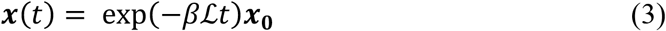

Let A be the observed FC network and Â be the predicted FC network from the ND model. We define the estimated FC of region *i* with all other regions at time *t* as the evolution on the graph of an initial configuration involving only region *i*, i.e. 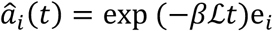 where e_*i*_ is the unit vector in the ith direction. If we collect all regions/unit vectors together, we obtain 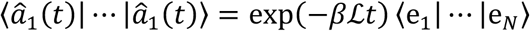, or

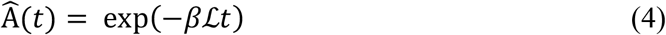

which gives the prediction for the observed FC matrix A. The accuracy of this prediction depends on *t* and *β.* We do not have an empirical value for *β*, so we absorb it into the estimation (by setting it to 1) and allowing *t* to vary. The special cases *Â*(0) = ***I*** and *Â*(∞) = *D*, where 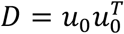 is the steady state solution (outer product of the eigenvector of 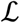 that has a corresponding eigenvalue of 0). Between those cases, a range of functional networks exists. The *t* that gives the most accurate predicted FC compared to the subject’s observed FC is called *ND model propagation time*, denoted *t_m_*. Specifically, ND model propagation time is the *t* that maximizes

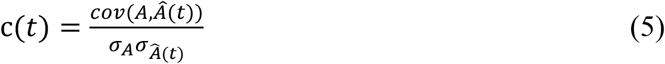

Here A and Â are vectorized versions of the matrices after excluding values on the diagonal. In summary, this procedure uses the ND model to estimate an individual’s FC from their SC and then identifies the *t* that gives the best agreement between the predicted and observed FC, which we call model propagation time. We understand model propagation time, which is unit-less and not related to actual time, as the measure of how much of the SC network is being used for the spread of functional activation as captured with the observed FC network.

### 2.5 Statistical Analysis

Changes in network metrics from 1 month to 6 months in the mTBI patients were calculated using a two-tailed, paired t-test (degrees of freedom = 25), while differences between mTBI (at both 1 month and 6 months) and healthy controls were assessed using an un-paired t-test (degrees of freedom = 58). QQ-plots of the connectome measures were used to verify normality of the connectome measures. P-values for all three sets of t-tests were corrected for multiple comparisons using Benjamini-Hochberg false discovery rate correction and assessed for significance using a threshold of α=0.05. Pearson’s correlation was calculated between the FC and SC at 1 month and 6 months, and between the change in network metrics between 1 and 6 months to evaluate the evolving relationship of FC and SC network metrics (degrees of freedom = 25). Two-tailed p-values were again Benjamini-Hochberg false discovery rate corrected and assessed for significance using a threshold of α=0.05.

For analysis of the relationship between recovery and connectome measures, we first used Principal Components Analysis (PCA) on the 25 subscores of attention (from ANT) and memory/learning (CVLT-II) on the concatenated data from all 27 subjects’ from both one and 6 months. The first principal component was calculated and taken to be a measure of overall cognition; differences between this measure at 1 and 6 months were compared using a two-tailed, paired t-test (degrees of freedom = 25) and assessed with a significance level of α=0.05. Once an overall measure of recovery was identified, Partial Least Squares Regression (PLSR), a regression technique that can accommodate correlated input variables, was used to model the relationship between changes in global connectome measures and changes in overall cognitive function. Specifically, we estimated change in overall cognitive recovery (*ΔPCA = PCA_FU_ — PCA_BL_*) from the various demographics and connectome metrics. The input variables included in the model were those of age, gender, change in ND model propagation time and change in the FC and SC global network metrics of average node degree, characteristic path length, global and local efficiency, clustering coefficient, modularity, transitivity, mean first passage time, mean navigation time on the structural connectome and mean navigation time based on Euclidean distance (*Am = m_FU_ — m_BL_*) that had trends for correlations (p<0.10 uncorrected) with the change in overall cognition. We performed this step as to not include any variables that were clearly not related to change in overall cognition. We used a nested cross-validation procedure to select and fit the models and perform predictions. The outer loop consisted of leave-one-out cross validation; each observation was held out in turn and the following procedure performed on the remaining training data to select and fit the model. First, the number of components in the PLSR model was chosen as the one that most frequently minimized the Predicted Residual Sum of Squares [Krishnan et al., 2011], calculated via k-fold (k=5) cross-validation with 50 Monte Carlo repetitions, over 1000 bootstrapped samples. Once the optimal number of components was identified, bootstrapping was again employed (with 10000 resamples having 50 Monte Carlo repetitions each) using the entire training data set to calculate the regression coefficients and the bias corrected and accelerated 95% confidence intervals [Efron, 1987]. The mean of the regression coefficients over the bootstrapped samples was then used to make a prediction for the single set of hold out test data. We assessed model performance by calculating the coefficient of determination (R^2^ = 1-SS_res_/SS_tot_) of the predicted values from the leave-one-out cross-validation. The data that support the findings of this study are available from the corresponding author upon reasonable request.

## 3.0 Results

### 3.1 Post-mTBI Cognitive Recovery

Figure 2 shows the first PCA component’s coefficients for the 9 subscores of the ANT and the 16 subscores (standardized) of the CVLT-II over the 27 subjects’ data from one and 6 months. Black lines indicate the 95% confidence intervals of each subscore’s PCA coefficient, calculated via bootstrapping; the weights used in the analysis are all well within the CIs. The first PCA component explained 48% of the variance, while the second and third components explained only 16% and 8%, respectively. The red bars signify that lower scores on that sub-test indicate better function while blue bars signify that higher scores on that sub-test indicate better function. All of the red bars have negative PCA coefficients (except for CVLT-II “repetitions”, which has a relatively small positive coefficient) and all blue bars are positive. This indicates that positive values of the PCA component indicate better cognitive scores, and increases over time indicate improvements in cognition. A paired t-test showed a significant increase in the PCA measure of overall cognition from one to 6 months (t_25_ = −2.15, p = 0.04), see Figure 3. In a secondary analysis (Supplementary Analysis S1), we included all available neuropsychological data from all time points and re-performed the PCA analysis (see Supplementary Figure S1), which was almost identical to the PCA results in Figure 2. Supplementary Figure S2 also shows violin plots of overall cognition from 1 week to 1 month and 6 months to 12 months, neither of which showed significant changes over time, supporting our hypothesis that most recovery would occur between 1 month and 6 months.

**Figure 2:**
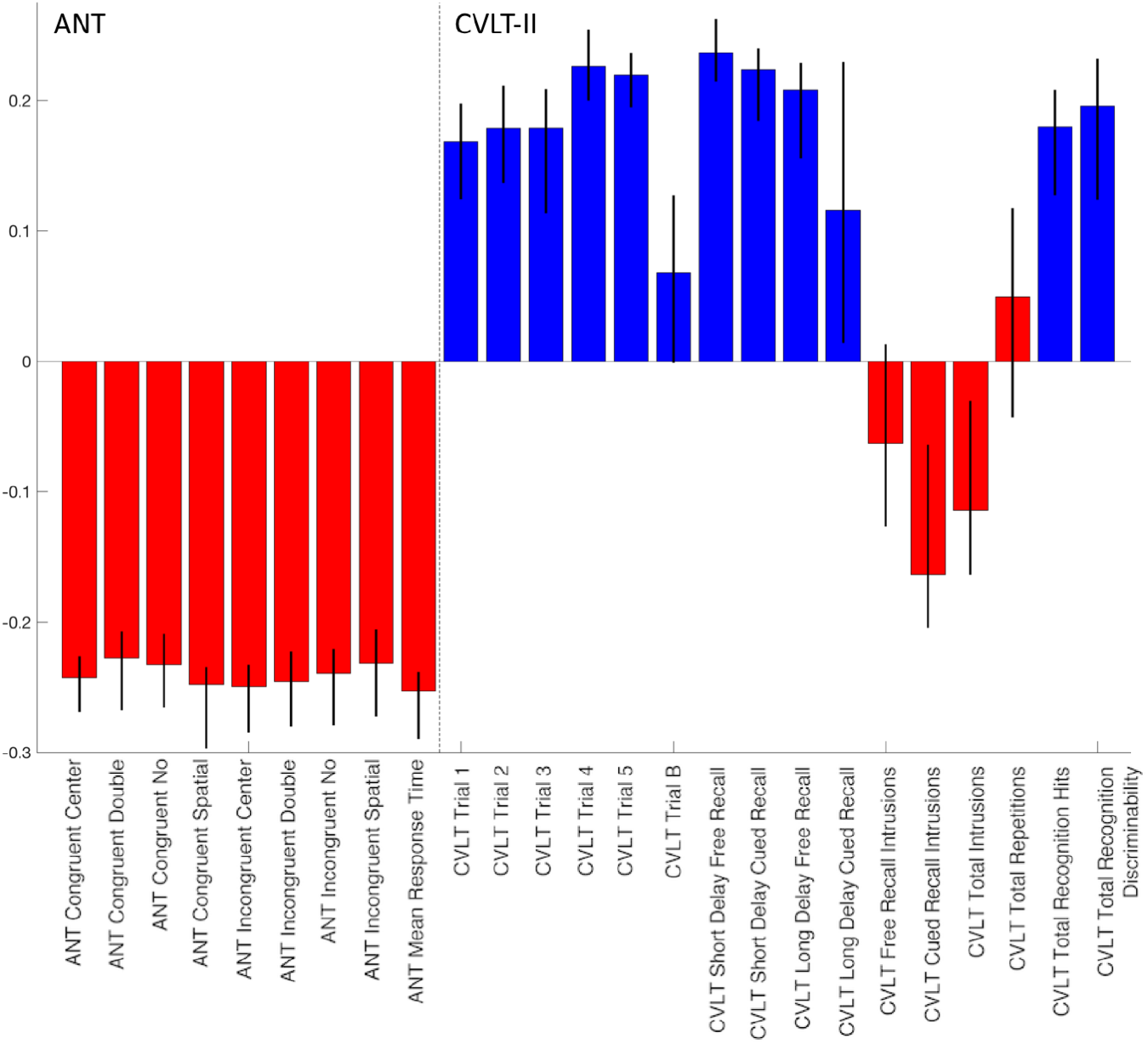
PCA coefficients of overall recovery. The coefficients of the first component of the PCA analysis using 27 subjects’ data from one and 6 months, with 95% bootstrapped confidence intervals superimposed as black lines. ANT scores are on the left and CVLT-II scores are on the right. Red indicates that sub-score’s values are smaller = better and blue indicates that sub-score’s values are higher = better.

**Figure 3:**
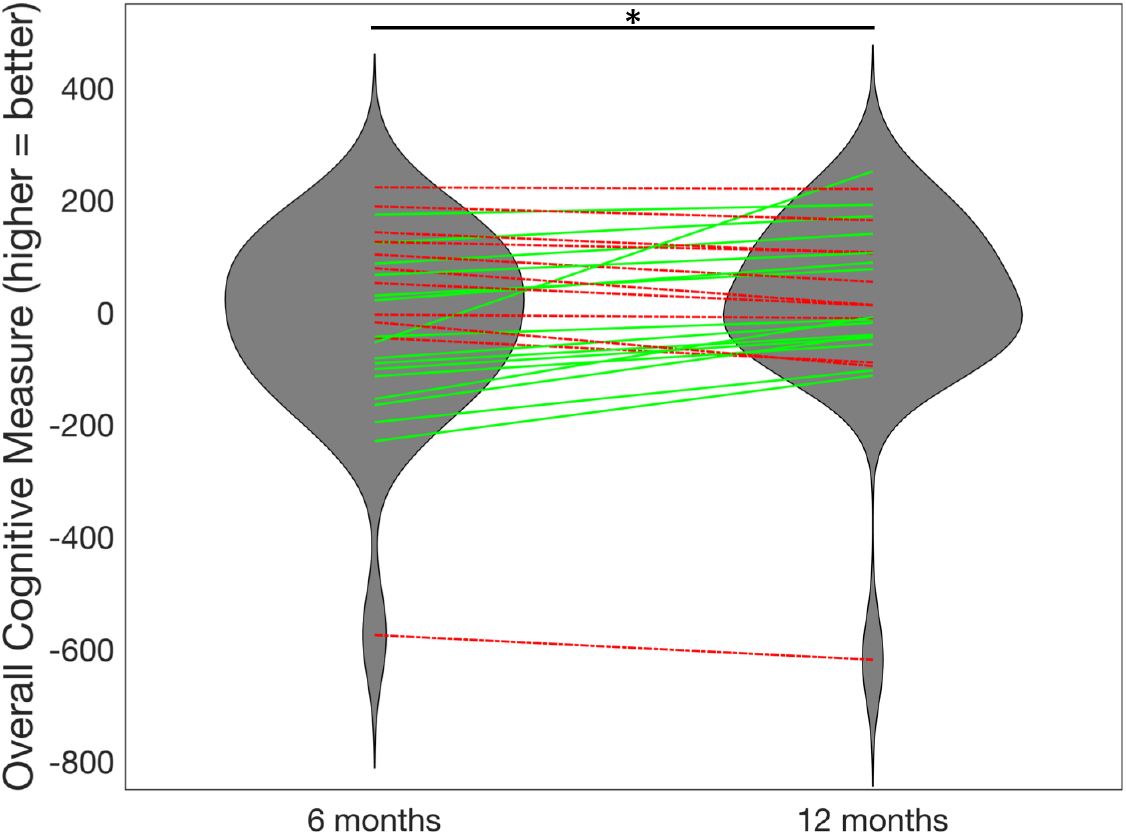
Longitudinal improvement in overall cognition. The violin plots describe the overall cognitive scores (first PCA component) at 1 months and 6 months, with lines indicating individuals (N = 27). Individuals with increases in cognition from 1 month to 6 months are plotted in green, solid lines while decreases in cognition are plotted in red, dashed lines. *Significant improvement in overall cognitive scores from 1 month to 6 months (p<0.05).

### 3.2 Network Diffusion Model Prediction

Figure 4A and 4B show the trajectories of the correlation between the observed FC and the ND model’s predicted FC over model time, with the individual’s maximum correlation (i.e. ND model propagation time) indicated with a red point, for the TBI and control subjects. Histograms of the final correlations between observed and predicted FC are given in Figure 4B (TBI subjects) and 4C (control subjects). Supplemental Video S1 (control subject) and Supplemental Video S2 (TBI subject) show a time-lapsed video of the observed FC, observed SC, the ND model’s predicted FC and the correlation between observed and predicted FC over model time for a particular individual. The point at which the correlation curve reaches its maximum is the final ND model propagation time. Correlations between the ND model’s predicted FC and observed FC range between 0.17 and 0.30, which is on par with the two previous studies using the ND model [Abdelnour et al., 2014; Kuceyeski et al., 2016], and similar to other studies of models predicting FC from SC [Falcon et al., 2016]. For comparison, we correlated the SC network with the observed FC and in all TBI and control subjects the ND model prediction correlations with observed FC were higher (see Supplementary Figure S3). In fact, a paired t-test of the sets of correlations revealed that the ND model predictions had significantly higher correlations with observed FC than a model based only on SC (t = 42.3, p ≈ 0).

**Figure 4:**
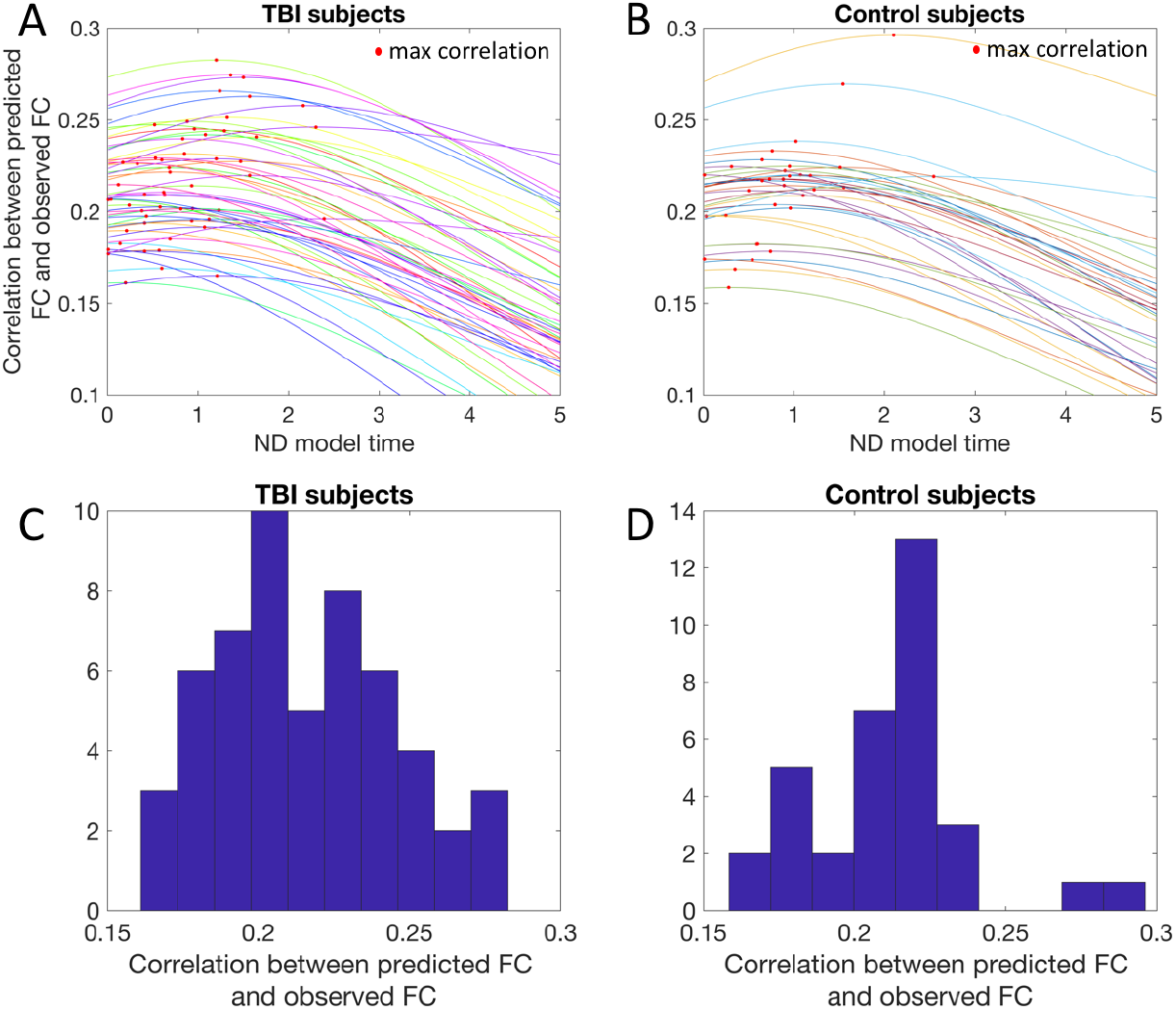
Network Diffusion (ND) model prediction. Panels A (TBI subjects) and B (controls) show the trajectories of the correlation between the observed functional connectivity and the ND model’s predicted functional connectivity over ND model time, with the individual’s maximum correlation (i.e. ND model propagation time) indicated with a red point. Panels C (TBI subjects) and D (controls) show the histograms of the final correlations between observed functional connectivity and the ND model’s predicted functional connectivity.

### 3.3 Predicting overall recovery from changes in global network metrics

One subject had an improvement in cognitive function between 1 and 6 months that was more than 1.5 times the inter-quartile range above the third quartile and therefore was excluded from the analyses. The final 26 mTBI patients with data from 1 and 6 months had demographics (29.4±8.0 years of age, 20 male) that were not significantly different than the entire mTBI population or the control group. There were no significant differences in any of the global SC and FC network metrics from 1 to 6 months and no significant differences at either time point when compared to healthy controls (p > 0.05, corrected for all t-tests), see Supplementary Table S1 for details. Correlations between demographics, change in graph network metrics and change in overall cognition are listed in Supplementary Table S2. The variables that showed trends for a relationship with change in cognition (uncorrected p<0.10) were change in ND model propagation time, SC characteristic path length, SC global efficiency, SC small-worldness, FC degree, FC characteristic path length, FC modularity and FC ratio of between to within module connection strength. These variables were then used as inputs to the PLSR models. All 26 PLSR models (over the leave-one-out cross-validation outer loop) included one component only. The predicted values versus observed values are provided in Figure 5A (hold-out coefficient of determination R^2^ = 0.27). Because we had one PLSR model for each of the leave-one-out iterations (which itself is the mean over the 10000 bootstrapped samples), we report the mean regression coefficient over all 26 models and list the number of times the confidence interval for the regression coefficient did not include 0 in Table 1. Violin plots of the regression coefficients for each input variable over each PLSR model is given in Figure 5B, where color indicates coefficient sign (red = positive, blue = negative).

**Figure 5:**
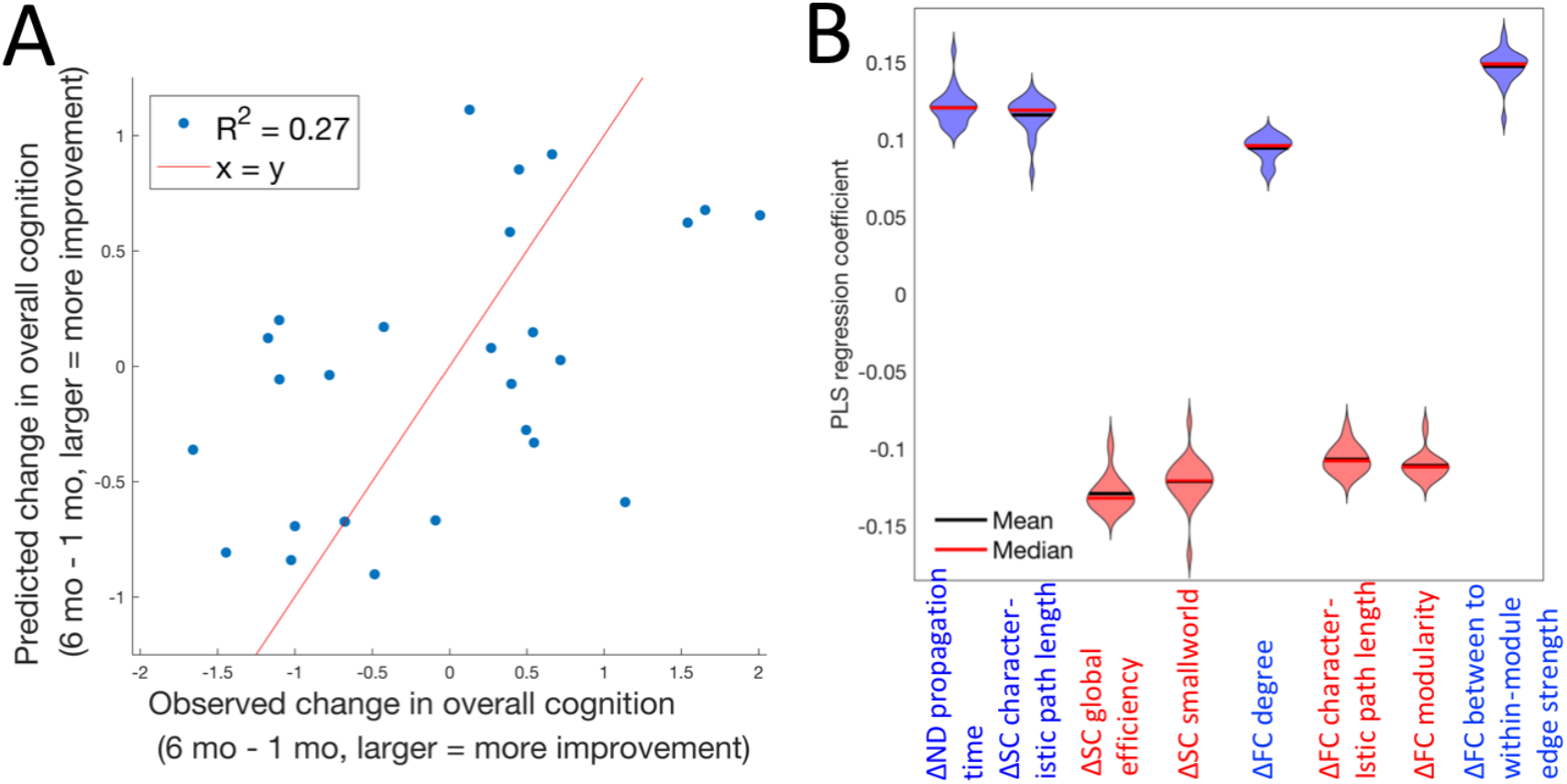
PLSR model of change in global network metrics predicting recovery. A) Observed versus predicted change in overall cognition post-mild TBI (the line of identity, x = y, is in red for reference), in normalized principal component units. The corresponding coefficient of determination (R^2^) is reported in the upper left corner. B) Violin plots indicating the shape of the distribution of coefficients over the 26 leave-one-out PLSR models predicting overall cognitive recovery post-mild TBI from changes in various FC and SC global network metrics. Red lines indicate the median while the blue lines indicate the mean. Violin plots are colored red or blue to indicate direction of relationship with recovery, where red indicates negative coefficients (decreases over time related to better recovery) while blue indicates positive coefficients (increases over time related to better recovery).

## 4.0 Discussion

We did not detect any significant FC or SC network metric differences between mTBI and healthy control groups, or any significant changes from 1 month to 6 months within the mTBI group. However, we observe that increases from 1 month to 6 months post-injury in network diffusion model propagation time, a measure of the relationship between FC and SC, was one of the predictors of improvements in overall cognitive function. ND model propagation time can be interpreted as a measure of the amount of SC that is utilized for the spread of functional activation that is captured via the FC network. The model coefficient was positive, indicating that increases in this value were associated with more improvement in cognitive functioning. ND model propagation time is inherently multimodal, capturing longitudinal changes in both SC and FC. In a post-hoc analysis, we investigated the Pearson correlation between the changes in the various global FC and SC network metrics. Supplementary Figure S4 shows that longitudinal increases in ND model propagation time had (uncorrected) correlations with, among other measures, increased SC characteristic path length (r = 0.44, p = 0.02) and decreased FC characteristic path length (r = −0.46, p = 0.02), both of which were also related to better recovery. It is important to note that the individual FC and SC metrics included in the PLSR model and ND model propagation time have similar contributions to overall cognitive recovery, as suggested by the magnitude of the PLSR coefficients in Table 1. ND model propagation time is a model-based, higher-order measure of both SC and FC and thus likely has less precision than the individual FC or SC metrics. The moderate correlation between FC metrics, SC network metrics and ND model propagation time also may dampen the contribution of any metric over another.

We conjecture that ND model propagation time captures the interplay between two biological post-TBI mechanisms, one of longitudinal segregation of the structural connectome and integration of the functional connectome (see Figure 1). The first, continued degeneration of white matter, is also evidenced by increased measures of segregation in the structural connectome in patients with better recovery, i.e. increased characteristic path length and decreased efficiency. This mechanism of remote degeneration may be more prevalent in the TBI patients with worse injuries that have more baseline impairment and thus more room for improvement. While some Diffusion Tensor Imaging (DTI) studies of longitudinal white matter changes post-TBI have shown continued white matter degeneration [Mayer et al., 2011; Niogi and Mukherjee, 2010; Palacios et al., 2018], others have shown no change or elevations in summary statistics of white matter microstructural integrity [Eierud et al., 2014]. These discrepancies may be due to the limitations of DTI in that it assumes Gaussian diffusion and the single modeled tensor used to calculate summary statistics does not accurately represent underlying complex white matter architecture, i.e. crossing and kissing fibers [Jones and Cercignani, 2010]. The second post-TBI recovery mechanism we conjecture is captured by increased ND model propagation time is one of neuroplasticity, in which increased integration of the functional connectome, also evidenced by decreased FC characteristic path length and modularity and increased ratio of between to within-module mean edge strength, may be a compensatory mechanism in response to the initial injury and/or continued white matter degeneration.

To our knowledge, this study is one of the first to quantify the longitudinal relationship between the functional and structural connectomes in the context of global cerebral reorganization after traumatic brain injury. It can be appreciated from Figure 3 that the amount of recovery in this mild TBI population is relatively small, although significant. Thus, the change in cognition has a relatively weak signal-to-noise ratio and may be quite difficult to predict. The fact that we were able to explain over 25% of this minor change with our global network measures in a moderate-sized cohort is an argument for the strength of our findings. We hypothesize that a longitudinal study of moderate to severe TBI would reveal similar relationships. In fact, our recent publication [Kuceyeski et al., 2016] performed cross-sectional analyses in severe brain injury patients (most of which had TBI etiology) to show that ND model propagation time was positively correlated with better measures of recovery of consciousness while other global measures of FC and SC were not. There, we also simulated injury and post-injury recovery using healthy connectomes to explore their impact on ND model propagation time. We simulated injury by removing random entries in the SC network and reducing the magnitude of FC in those same entries by a varying percent (25-100%). We simulated recovery by removing random entries in the SC and leaving the FC of those region-pairs intact. Using these simulated networks, we showed that ND model propagation time had decreases in injury and increases in recovery that were proportional to the amount of injury and recovery (see Fig 6 of Kuceyeski *et al.*, 2016), which provides further support for our claim that increased ND model propagation time may capture multi-modal post-injury mechanisms of increased structural segregation and increased functional integration. As compared to this previous study, our current study is 1) longitudinal (the previous paper was cross-sectional), 2) in patients with TBI (the previous paper’s cohort had varying etiology of injury), 3) in patients with mild injury (the previous paper’s cohort had severe injury) and 4) based on cognitive measures ANT/CVLT-II that are sensitive to much lesser degrees of brain dysfunction (previously we used the Coma Recovery Scale – Revised measure that assesses consciousness). Yet, we still observe that ND model propagation time increases with measures of recovery. This robust finding lends confidence that our observations in mild TBI are not a result of chance and that ND model propagation time is indeed capturing some mechanism of global network-level neuroplasticity that is related to recovery after injury.

### 4.1 Comparison to previous work

Studies that have investigated global network metrics in mTBI have found mixed results (see Caeyenberghs et al., [2017] for a review), probably due in part to the heterogeneity of the disorder and the populations studied in addition to the complicated relationship between FC/SC and impairment and recovery. Many studies have found no group-wise differences when comparing mTBI and healthy controls’ global network metrics or when comparing longitudinal changes in the mild TBI group [Hillary et al., 2014; van der Horn et al., 2017b], which agree with our findings here. While van der Horn et al., [2017b] found no differences in SC global network metrics between TBI and controls, lower global and mean local efficiency were found in the SC networks of TBI patients without post-traumatic complaints when compared to those with post-traumatic complaints, which is the crosssectional analog of our longitudinal result that decreased SC global efficiency over time was related to better cognitive recovery. In contrast, other studies have found group-wise differences between mTBI and healthy controls’ global network measures. For example, Pandit et al. [2013] found reduced overall FC, longer characteristic path lengths and reduced efficiency in mTBI patients versus controls, while Nakamura et al., [2009] showed lower small world indices in the resting-state FC network. Another study showed that moderate to severe TBI patients had lower global SC network efficiency than normal controls and that lower global efficiency was also correlated with worse scores on an executive function task [Caeyenberghs et al., 2014]. The discrepancy between their findings and our results could be due to the difference in populations, i.e. moderate to severe versus mild TBI. Kim et al., [2014] showed no differences in transitivity and modularity between mTBI and controls, but showed longer SC characteristic path lengths were moderately correlated with worse performance on executive function and verbal learning tasks in the mild TBI group. However, multiple comparisons corrections were not performed in this cross-sectional, preliminary study.

Only a few studies have investigated the interplay between FC and SC changes in recovery after mTBI. One such study showed TBI patients with more SC injury had less FC in the default mode network [Sharp et al., 2011]. They also showed that higher resting-state FC in the posterior cingulate cortex was correlated with more efficient response speeds. Another analysis of task-based FC and SC networks in mTBI showed that there were no correlations between FC and SC network metrics, no differences in the SC network metrics and significant increases in FC strength in patients versus controls [Caeyenberghs et al., 2013]. In another study of chronic TBI patients, Palacios et al., [2013] found increases in FC in frontal areas compared to controls that was positively associated with better cognitive outcomes and negatively associated with a measure of SC. They concluded that altered SC between brain regions could be partly compensated for by increased FC. This result is also supported by Bonnelle et al., [2012], wherein the authors showed a failure of default mode network deactivation was associated with impairment after TBI, and that this abnormal default mode network FC could be predicted by the amount of SC disruption in salience network regions, specifically right anterior insula to pre-SMA and dorsal anterior cingulate. These cross-sectional studies are compatible with our current findings of longitudinal increases in ND model propagation, increases in FC degree, decreases in FC characteristic path length, increases in SC characteristic path length and decreases in SC efficiency in recovery. Also in line with these findings are the moderate associations we observed between measures of SC network segregation and FC network integration over time (Fig S4), e.g. SC global efficiency and FC degree (r = −0.41, p = 0.04) and SC characteristic path length and FC global efficiency (r = 0.41, p = 0.04).

### 4.2 Post-hoc regional connectivity analysis

In a post-hoc analysis, we calculated regional node strength (sum of all connections per node) for both FC and SC and calculated the differences in the mTBI patients versus the controls and the change over time in the mTBI subjects. To improve the signal to noise ratio of node strength and reduce the number of comparisons/model inputs, we first averaged node strength over the left and right hemispheres and performed the analyses over 43 ROIs. We calculated node strength over the original FC, which included positive and negative values. Supplemental Table S3 and S4 and Supplemental Figure S5 give the t-statistics of the three group-wise comparisons for functional and structural node strength. None of the t-statistics had corrected p-values that were significant, likely due to the small sample size and large number of comparisons. Here we only discuss trends in the group comparisons. In general, we see more regions with greater FC node strength and more regions with weaker SC node strength in the mTBI population compared to controls. This weaker SC node strength was particularly evident in the cerebellum; the temporal pole displayed a trend for stronger SC node strength in mTBI compared to controls. We see trends for increased SC node strength over time in the mTBI subjects in the caudal middle frontal and precuneus, while weaker trends exist for decreased SC node strength in the paracentral gyrus and posterior cingulate. We see especially large FC node strength in the mTBI patients compared to controls in the caudal anterior cingulate, while FC appeared to be lower in mTBI in the inferior temporal region. FC node strength tended to increase over time in the mTBI population in the fusiform gyrus and decrease over time in the thalamus, inferior parietal, caudal middle frontal and precuneus regions.

Finally, to quantify the relationship between change in regional node strength and recovery, we built PLSR models in the same manner described above to predict 1) overall recovery, 2) ANT’s mean reaction time (attention) and 3) CVLT-II’s mean of trials 1-5 (working memory) from change in FC and SC node strength over time. Figure 6 shows the scatter plots of predicted versus observed recovery measures for the three different outcome measures, with violin plots of the regression coefficients in the bottom panel. Figure 7 visualizes the regional regression coefficients for the models of each cognitive recovery measure, with blue indicating increases in that region’s functional or structural node strength were related to better recovery and red indicating that decreases in node degree were related to better recovery. The model of overall cognition and ANT’s mean reaction time are almost identical, which is likely due to the PCA component having such a large coefficient for this cognitive subscore. We see that increased SC node strength in the superior temporal sulcus and decreased SC node strength in sensory-motor regions, including precentral and postcentral gyrus, was related to better overall cognitive/attention recovery. Increases in many regions’ FC node strength, including the thalamus, hippocampus, inferior temporal, pars triangularis, supramarginal gyrus and insula were related to better overall cognitive/attention recovery, while decreases in FC node strength in the caudal anterior cingulate were related to better overall cognitive/attention recovery. The regions whose improved FC are associated with better cognitive recovery consist of well-known connectome hubs, including subcortical (thalamus) and archicortical (hippocampus), as well as key components of the default mode network (supramarginal gyrus and inferior temporal gyrus) and salience network (insula). Hence, these particular regions are postulated to be the greatest beneficiaries of increased functional integration in recovery.

**Figure 6:**
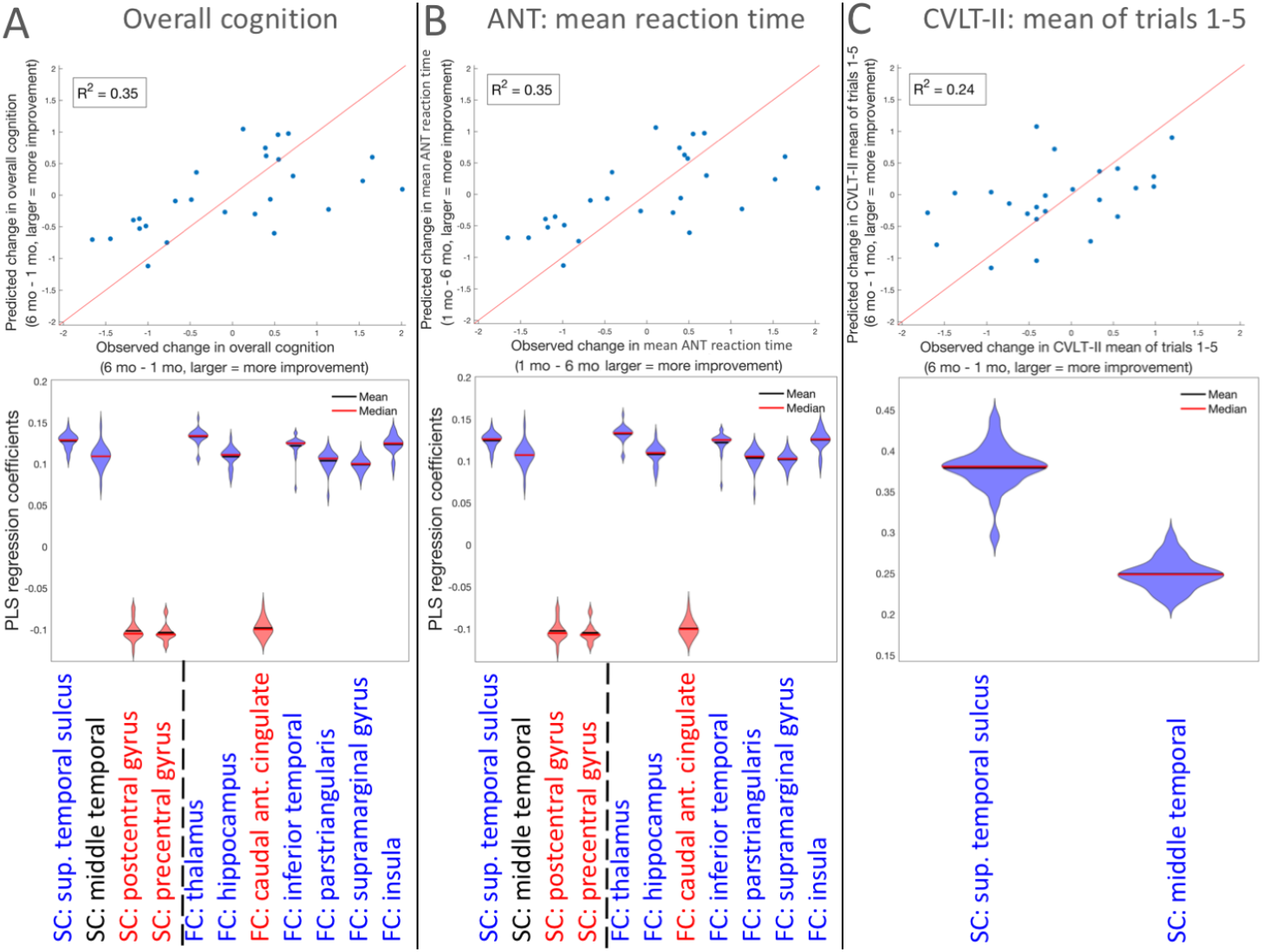
PLSR model of change in regional node strength predicting recovery. PLSR model results for change in regional node strength predicting the change in A) overall cognition (PCA), B) ANT’s mean reaction time and C) CVLT-II’s mean over Trials 1-5. Top panels are scatter plots of the observed versus predicted cognitive function (line of identity, x = y, is in red for reference). Bottom panels are violin plots for the 26 coefficients created in each of the leave-one-out models. Violin plots are colored red or blue to indicate direction of relationship with recovery, where red indicates negative coefficients (decreases over time related to better recovery) while blue indicates positive coefficients (increases over time related to better recovery).

The role of thalamus in attention and recovery post-TBI has been well-documented [Schiff et al., 2007]. The thalamus and the caudal anterior cingulate are also part of the anterior meso-circuit, which is hypothesized to play an integral part in attention recovery after TBI [Fridman et al., 2014; Schiff, 2010]. Interestingly, mTBI patients showed a non-significant trend toward greater caudal anterior cingulate FC than controls at 1 month post-injury (see Supplemental Table S4), and those patients with an interval decrease in FC of that region by 6 months after mTBI experienced better cognitive recovery, especially for visuospatial attention as measured by the ANT (Figure 6B). This agrees with the known role of the anterior cingulate in attentional focus and cognitive control, with elevated functional activation early after TBI and progressively reduced activation associated with improving task performance [Cazalis et al., 2011; Scheibel, 2017]. Increases in SC node strength in temporal regions (superior temporal sulcus and middle temporal gyrus) were related to better recovery of verbal memory, which is in agreement with the known role of the temporal lobe in memory and language function.

**Figure 7:**
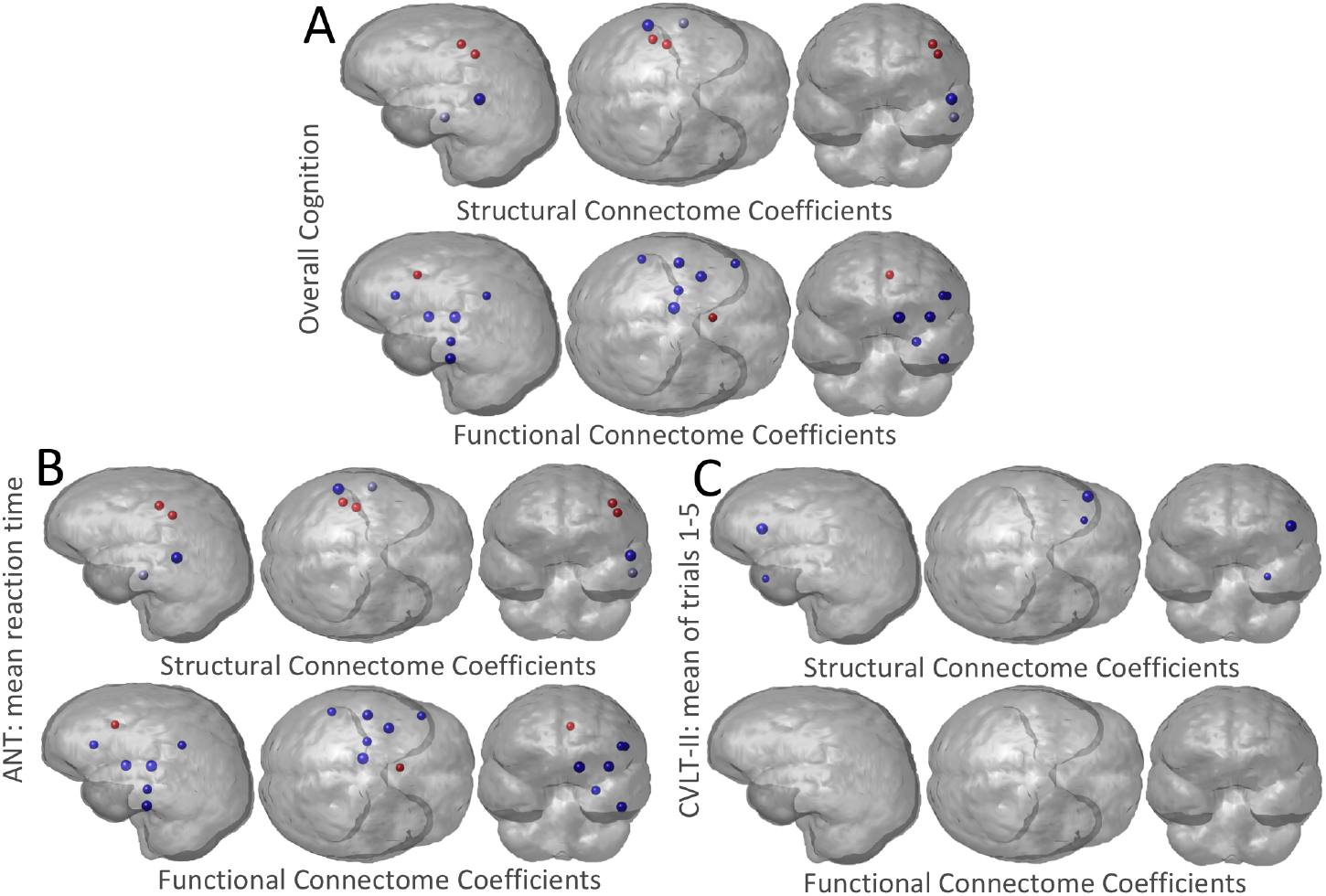
Glassbrain visualization of the regional PLSR coefficients. Three PLSR models were constructed to predict change in A) overall cognition, B) ANT’s mean reaction time and C) CVLT-II’s mean of trails 1-5, from change in FC and SC node strength over time in the TBI population (6 months – 1 month). The PLSR coefficient for each region (node strength was averaged over hemispheres) is represented by a sphere, where the size is proportional to the magnitude of the PLSR regression coefficient. Blue regions indicate those regions where increases in node strength were related to better recovery, while red indicates those regions where decreases in node strength were related to worse recovery. Results shown for left cerebral hemisphere only due to the combination of left/right region pairs in the analysis.

### 4.3 Limitations

A limitation of the current work is the relatively small sample size. To combat the effects of the small sample size, we performed leave-one-out cross-validation and bootstrapping for model selection and inference. There are also some limitations in the data processing. We did not have fieldmaps with which to perform EPI distortion correction for dMRI. However, the degree of anatomical distortion was low in this data due to relatively strong gradients with a short TE of 63 ms, as well as the high spatial resolution of 1.8-mm isotropic voxels. Tractography algorithms have issues reconstructing fibers that are crossing and kissing – here we use multi-tensor fitting of the dMRI to minimize this issue. The FC networks represent correlations of time-series and not physical connections. Therefore, although they are widely used in the literature [Wang et al., 2010], interpretation of some of the network metrics like characteristic path length and efficiency may not be as straightforward as the same measures in structural connectivity networks. Interpretation is particularly challenging when considering negative entries, which is why we remove them in our current analyses. Finally, previous studies in mTBI have shown correlations between attention, memory and depression measures and connectivity metrics in particular brain regions [Bonnelle et al., 2011; Hampson et al., 2006; van der Horn et al., 2017a; Sharp et al., 2011]. We focused here on global connectomic measures since our sample size was not large and we wanted to minimize the effect of heterogeneity of injury patterns and reduce the number of statistical tests/input variables. We believe global network metrics would be more robust to the heterogeneity in patient injury patterns and the global nature of diffuse axonal injury. Therefore, we only perform a post-hoc analysis of regional abnormality/changes in FC and SC node strength, and interpret the findings with caution. We believe that investigating functional reorganization on a regional basis would be an excellent area of research, and we plan to do this with larger sets of data.

### 4.4 Conclusions and future work

Gaining a clear picture of the mechanism driving cerebral reorganization for a particular individual’s pattern of brain injury will enable the development of biomarkers for more accurate prognoses and development of personalized treatment plans using a Precision Medicine framework [Collins and Varmus, 2015]. These personalized treatments could be based on cognitive or physical therapeutic approaches, or they could be physiological, e.g. non-invasive brain stimulation. Non-invasive brain stimulation has been shown to modify the brain’s FC networks to boost recovery from stroke, depression and mTBI [Demirtas-Tatlidede et al., 2013; Grefkes and Fink, 2011]. It is not clear how neuromodulation techniques influence the brain, but increases in FC between regions have been shown in high-frequency repetitive transcranial magnetic stimulation (rTMS) [Thut and Pascual-Leone, 2010]. Currently, the choice of targets for brain stimulation is not well defined; many times it depends on population-level observations. If we can fully understand the mechanism of post-injury cerebral organization in terms of the structural and functional connectome relationship, then we may be able to identify region-pairs in a particular individual that, if functionally connected, would have the largest influence on improvements in attention and memory. We would be able to explicitly identify structural pathways that could be used to establish this functional connection. Such regions and pathways would then be optimal targets for rTMS. This method for personalized target selection could be applied in a variety of neurological disorders, improving recovery and quality of life for patients with a range of neurological diseases.

## Acknowledgements

This work was supported by an Anna-Maria and Stephen Kellen Foundation Junior Faculty Fellowship (AK) and the NIH [R01 NS060776 (PM), R21 NS104634-01 (AK), R01 NS102646-01A1 (AK), R01NS092802 and R01 EB022717 (AR)]. PM has received research support from GE Healthcare and serves on the Medical Advisory Board of the GE-NFL Head Health Initiative.

## Notes

#### Summary of Updates

Functional connectivity matrices were handled differently in the calculation of the graph theoretical measures - in the first version the absolute value was taken and in this version the negative entries are replaced with zero to allow easier interpretation. The hypothesized recovery mechanism that our results support has thus changed from functional rerouting to a dual network based mechanism wherein integration of the functional connectome and segregation of the structural connectome is related to better recovery (see new Figure 1). A few more graph theoretical measures to support this new theory were also investigated in this version and additional analyses of the ND model prediction accuracy were performed.

