## Supplemental Data 1 for "Longitudinal increases in structural connectome segregation and functional connectome integration are associated with better recovery after mild TBI"

**Supplementary Material**

*Supplemental Analysis A1: Overall cognitive recovery at 1 week, 1 month, 6 months and 12 months*

There were 138 datasets with complete neuropsychological data available out of 204 (4x51) possible: 26 from 1 week, 39 from 1 month, 37 from 6 months and 36 from 1 year. We put all of the available neuropsychological data into a PCA analysis, and got coefficients for each of the 25 neuropsychological tests (see Supplementary Fig. 1) that were almost identical to those found in the analysis of the 27 subjects from 1 month and 6 months only (see Fig. 2 of the main paper). In fact, the overall cognitive scores derived from PCA on the entire dataset versus PCA on just the 27 subjects from 1 month and 6 months (the data included in the main analysis) was r = 1.0 (p ≈ 0).

There were 11 subjects with imaging and neuropsychological data from 1 week and 1 month, 27 subjects with both 1 month and 6 months and 18 subjects with both 6 months and 12 months. A paired t-test showed the largest sequential increase in the PCA measure of overall cognition occurred between 1 month and 6 months (t = -2.12, p = 0.043), compared to the change from 1 week to 1 month (t = -1.00, p = 0.33) or from 6 months to 1 year (t = -1.20, p = 0.24), see Supplementary Fig. 2 and Fig. 3 in the main paper.

|  | **TBI: BL vs. FU**  **t_25_ (p-value)** | **TBI: BL vs. HC**  **t_58_ (p-value)** | **TBI: FU vs. HC**  **t_58_ (p-value)** |
| --- | --- | --- | --- |
| ND model propagation time | -0.1991 (0.99) | -0.2356 (0.99) | -0.0834 (0.99) |
| SC node strength | -1.2246 (0.87) | -0.8168 (0.99) | -0.0436 (0.99) |
| SC characteristic path length | 0.3507 (0.99) | 0.3736 (0.99) | 0.1188 (0.99) |
| SC global efficiency | -1.4999 (0.69) | -0.6383 (0.99) | 0.1238 (0.99) |
| SC clustering coefficient | 1.7136 (0.61) | 2.0314 (0.61) | 0.4785 (0.99) |
| SC modularity | 1.9972 (0.61) | -0.5164 (0.99) | -2.8594 (0.44) |
| SC small-world metric | 0.8995 (0.99) | 1.0719 (0.95) | 0.2533 (0.99) |
| SC local efficiency | 1.6107 (0.64) | 1.9626 (0.61) | 0.5347 (0.99) |
| SC ratio of between to within module edge strength | -1.7253 (0.61) | 0.1311 (0.99) | 1.7839 (0.61) |
| SC transitivity | 1.6763 (0.61) | 2.4018 (0.61) | 0.9446 (0.99) |
| SC mean first passage time | -0.2341 (0.99) | 1.8635 (0.61) | 1.7797 (0.61) |
| SC mean navigation time | -1.1153 (0.95) | -0.7951 (0.99) | 0.0112 (0.99) |
| SC mean distance-navigation time | -0.9476 (0.99) | 0.7785 (0.99) | 1.7525 (0.61) |
| FC degree | -0.4192 (0.99) | 0.4775 (0.99) | 0.7337 (0.99) |
| FC characteristic path length | 0.6828 (0.99) | -0.3165 (0.99) | -0.7114 (0.99) |
| FC global efficiency | -0.4585 (0.99) | 0.2683 (0.99) | 0.5282 (0.99) |
| FC clustering coefficient | -0.0211 (0.99) | 0.5355 (0.99) | 0.5448 (0.99) |
| FC modularity | 0.0966 (0.99) | -1.0183 (0.98) | -1.2061 (0.87) |
| FC Small-world metric | -0.0387 (0.99) | -1.4382 (0.69) | -1.4968 (0.69) |
| FC local efficiency | 0.2022 (0.99) | 0.3469 (0.99) | 0.2008 (0.99) |
| FC ratio of between to within module edge strength | -0.2046 (0.99) | 1.3357 (0.78) | 1.7686 (0.61) |
| FC transitivity | 0.3972 (0.99) | 0.722 (0.99) | 0.4564 (0.99) |
| FC mean first passage time | 0.0299 (0.99) | 0.6144 (0.99) | 0.5366 (0.99) |
| FC mean navigation time | 0.322 (0.99) | 0.1003 (0.99) | -0.166 (0.99) |
| FC mean distance-navigation time | -0.4483 (0.99) | 0.6139 (0.99) | 1.0667 (0.95) |

**Supplementary Table S1:** Comparisons of FC and SC network metrics across time points within the traumatic brain injury (TBI) group at baseline (BL, 1 month) and follow-up (FU, 6 months) and between the two TBI time points and the healthy controls (HC).

|  | **Pearson’s r with overall cognition (uncorrected p-value, corrected p-value)** |
| --- | --- |
| Age | 0.0724 (0.73, 0.82) |
| Gender | -0.103 (0.62, 0.79) |
| **ND model propagation time** | **0.4332 (0.03, 0.17)*** |
| SC degree | -0.0884 (0.67, 0.82) |
| **SC characteristic path length** | **0.4306 (0.03, 0.17)*** |
| **SC global efficiency** | **-0.4787 (0.01, 0.17)*** |
| SC clustering coefficient | -0.1916 (0.35, 0.61) |
| SC modularity | 0.0756 (0.71, 0.82) |
| **SC small-world metric** | **-0.4114 (0.04, 0.17)*** |
| SC local efficiency | -0.2176 (0.29, 0.61) |
| SC ratio of between to within module edge strength | -0.0564 (0.78, 0.82) |
| SC transitivity | -0.3089 (0.12, 0.37) |
| SC mean first passage time | -0.1495 (0.47, 0.70) |
| SC mean navigation time | -0.2021 (0.32, 0.61) |
| SC mean distance-navigation time | -0.2041 (0.32, 0.61) |
| **FC degree** | **0.3577 (0.07, 0.25)*** |
| **FC characteristic path length** | **-0.3999 (0.04, 0.17)*** |
| FC global efficiency | 0.294 (0.14, 0.39) |
| FC clustering coefficient | 0.184 (0.37, 0.61) |
| **FC modularity** | **-0.4086 (0.04, 0.17)*** |
| FC Small-world metric | -0.2188 (0.28, 0.61) |
| FC local efficiency | 0.1148 (0.58, 0.78) |
| **FC ratio of between to within-module edge strength** | **0.5242 (0.01, 0.16)*** |
| FC transitivity | 0.1273 (0.54, 0.76) |
| FC mean first passage time | 0.055 (0.79, 0.82) |
| FC mean navigation time | -0.1782 (0.38, 0.61) |
| FC mean distance-navigation time | 0.0171 (0.93, 0.93) |

**Supplementary Table S2:** Correlation values (Pearson) between the change in each network metric and the change in overall cognitive scores. Uncorrected and corrected (using Benjamini-Hochberg) p-values are shown. *****Uncorrected p<0.10

|  | Controls vs.  1-mo. TBI  t-stat (p-value) | Controls vs.  6-mo. TBI  t-stat (p-value) | 1-mo. TBI vs 6-mo. TBI  t-stat (p-value) |
| --- | --- | --- | --- |
| Cerebellum | 3.1 (0.13) | 2.87 (0.12) | 0.24 (0.95) |
| Fusiform gyrus | 2.15 (0.36) | 1.47 (0.82) | 1.23 (0.77) |
| Cuneus | 2.06 (0.36) | 1.26 (0.82) | 1.28 (0.77) |
| Inferior temporal | 2 (0.36) | 1.13 (0.82) | 1.82 (0.74) |
| Caudal middle frontal | 1.64 (0.59) | 0.32 (0.92) | 2.68 (0.56) |
| Pericalcarine | 1.63 (0.59) | 1.02 (0.82) | 0.88 (0.77) |
| Transverse temporal | 1.56 (0.59) | 2.02 (0.69) | -0.83 (0.77) |
| Lateral occipital | 1.36 (0.65) | 0.78 (0.86) | 0.75 (0.77) |
| Parsorbitalis | 1.27 (0.65) | 0.89 (0.82) | 0.55 (0.87) |
| Parsopercularis | 1.24 (0.65) | 0.95 (0.82) | 0.45 (0.88) |
| Bank of the temporal sulcus | 1.22 (0.65) | 1.29 (0.82) | -0.07 (0.99) |
| Precentral | 1.17 (0.65) | 0.5 (0.92) | 0.74 (0.77) |
| Inferior parietal | 1.17 (0.65) | 0.63 (0.86) | 0.89 (0.77) |
| Precuneus | 1.1 (0.65) | -0.75 (0.86) | 2.17 (0.74) |
| Amygdala | 1.06 (0.65) | -0.29 (0.92) | 1.78 (0.74) |
| Supramarginal | 1.03 (0.65) | 1.13 (0.82) | -0.04 (0.99) |
| Isthmus cingulate | 0.97 (0.65) | 0.64 (0.86) | 0.39 (0.88) |
| Postcentral | 0.94 (0.65) | 0.15 (0.92) | 1 (0.77) |
| Middle temporal | 0.94 (0.65) | 0.98 (0.82) | 0.01 (0.99) |
| Thalamus | 0.91 (0.65) | 0.12 (0.92) | 0.83 (0.77) |
| Lingual gyrus | 0.8 (0.65) | 1.03 (0.82) | -0.47 (0.88) |
| Parahippocampal gyrus | 0.75 (0.67) | 0.45 (0.92) | 0.36 (0.88) |
| Superior Frontal | 0.57 (0.78) | 0.18 (0.92) | 0.76 (0.77) |
| Frontal pole | 0.57 (0.78) | -0.26 (0.92) | 1.22 (0.77) |
| Nucleus accumbens | 0.52 (0.78) | -1 (0.82) | 1.53 (0.77) |
| Hippocampus | 0.08 (0.97) | 0.22 (0.92) | -0.16 (0.96) |
| Hypothalamus | 0.07 (0.97) | 0.21 (0.92) | -0.16 (0.96) |
| Superior temporal | 0.03 (0.98) | 0.88 (0.82) | -0.92 (0.77) |
| Entorhinal | -0.13 (0.97) | -0.38 (0.92) | 0.24 (0.95) |
| Parstriangularis | -0.15 (0.97) | -0.05 (0.96) | -0.14 (0.96) |
| Medial orbitofrontal | -0.22 (0.94) | -0.93 (0.82) | 0.79 (0.77) |
| Globus Pallidus | -0.26 (0.93) | -0.98 (0.82) | 0.77 (0.77) |
| Lateral orbitofrontal | -0.29 (0.92) | 0.17 (0.92) | -0.54 (0.87) |
| Caudate | -0.42 (0.83) | -0.67 (0.86) | 0.36 (0.88) |
| Superior parietal | -0.5 (0.78) | -1.26 (0.82) | 0.95 (0.77) |
| Rostral middle frontal | -0.54 (0.78) | 0.15 (0.92) | -0.98 (0.77) |
| Rostral anterior cingulate | -0.8 (0.65) | -1.25 (0.82) | 0.64 (0.84) |
| Putamen | -0.84 (0.65) | -1.54 (0.82) | 1.28 (0.77) |
| Paracentral | -0.88 (0.65) | 0.53 (0.91) | -1.99 (0.74) |
| Caudal anterior cingulate | -1.05 (0.65) | -0.26 (0.92) | -0.77 (0.77) |
| Insula | -1.12 (0.65) | -0.62 (0.86) | -0.52 (0.87) |
| Posterior cingulate | -2.08 (0.36) | -0.63 (0.86) | -1.47 (0.77) |
| Temporalpole | -2.12 (0.36) | -3.33 (0.06) | 1.55 (0.77) |

**Supplementary Table S3:** Results of the unpaired group-wise t-tests of structural connectome node strength (averaged over the right and left homologues) between controls and TBI at 1 and 6 months and between TBI patients at 1 and 6 months.

|  | Controls vs.  1-mo. TBI  t-stat (p-value) | Controls vs.  6-mo. TBI  t-stat (p-value) | 1-mo. TBI vs. 6-mo. TBI  t-stat (p-value) |
| --- | --- | --- | --- |
| Inferior temporal | 2.43 (0.4) | 1.82 (0.64) | 0.87 (0.74) |
| Superior parietal | 1.83 (0.67) | 0.95 (0.72) | 1.15 (0.7) |
| Fusiform gyrus | 1.47 (0.69) | -0.5 (0.81) | 2.48 (0.49) |
| Parsopercularis | 1.05 (0.69) | 0.13 (0.93) | 0.89 (0.74) |
| Lingual gyrus | 0.9 (0.69) | -0.2 (0.93) | 1.04 (0.7) |
| Precentral | 0.84 (0.69) | -0.52 (0.81) | 1.4 (0.7) |
| Postcentral | 0.82 (0.69) | -0.42 (0.85) | 1.26 (0.7) |
| Parahippocampal gyrus | 0.75 (0.69) | 0.28 (0.91) | 0.42 (0.94) |
| Globus Pallidus | 0.73 (0.69) | -0.12 (0.93) | 0.79 (0.78) |
| Parsorbitalis | 0.68 (0.69) | -0.15 (0.93) | 0.91 (0.74) |
| Paracentral | 0.5 (0.76) | -0.53 (0.81) | 1.09 (0.7) |
| Parstriangularis | 0.41 (0.81) | -0.63 (0.81) | 1.05 (0.7) |
| Supramarginal | 0.4 (0.81) | -1.27 (0.72) | 1.65 (0.68) |
| Middle temporal | 0.24 (0.89) | 0.72 (0.76) | -0.49 (0.94) |
| Inferior parietal | 0.03 (0.97) | 2.2 (0.34) | -2.11 (0.49) |
| Transverse temporal | -0.04 (0.97) | -0.84 (0.72) | 1.13 (0.7) |
| Frontal pole | -0.05 (0.97) | -0.27 (0.91) | 0.24 (1) |
| Putamen | -0.19 (0.92) | -0.97 (0.72) | 1.11 (0.7) |
| Entorhinal | -0.33 (0.84) | -0.52 (0.81) | 0.23 (1) |
| Pericalcarine | -0.56 (0.73) | -0.6 (0.81) | 0.02 (1) |
| Caudal middle frontal | -0.61 (0.71) | 1.15 (0.72) | -2.08 (0.49) |
| Amygdala | -0.63 (0.71) | -0.93 (0.72) | 0.46 (0.94) |
| Caudate | -0.69 (0.69) | 0.05 (0.96) | -0.99 (0.71) |
| Bank of the superior temporal sulcus | -0.71 (0.69) | -1.3 (0.72) | 0.67 (0.85) |
| Hypothalamus | -0.71 (0.69) | 1.19 (0.72) | -1.69 (0.68) |
| Cerebellum | -0.72 (0.69) | -0.72 (0.76) | 0.01 (1) |
| Lateral occipital | -0.75 (0.69) | -1.38 (0.72) | 0.6 (0.88) |
| Cuneus | -0.9 (0.69) | -0.93 (0.72) | 0.05 (1) |
| Lateral orbitofrontal | -0.92 (0.69) | -0.91 (0.72) | -0.09 (1) |
| Rostral middle frontal | -0.94 (0.69) | -0.83 (0.72) | -0.2 (1) |
| Hippocampus | -0.98 (0.69) | -0.82 (0.72) | -0.06 (1) |
| Posterior cingulate | -1.01 (0.69) | -1.63 (0.72) | 0.66 (0.85) |
| Superior Frontal | -1.01 (0.69) | -1.06 (0.72) | 0 (1) |
| Thalamus | -1.1 (0.69) | 1.17 (0.72) | -2.23 (0.49) |
| Superior temporal | -1.1 (0.69) | -2.23 (0.34) | 1.19 (0.7) |
| Nucleus accumbens | -1.14 (0.69) | -0.94 (0.72) | -0.42 (0.94) |
| Rostral anterior cingulate | -1.15 (0.69) | -1.22 (0.72) | 0.12 (1) |
| Insula | -1.43 (0.69) | -2.52 (0.31) | 1.4 (0.7) |
| Temporalpole | -1.6 (0.69) | -1.27 (0.72) | -0.26 (1) |
| Medial orbitofrontal | -1.7 (0.67) | -1.02 (0.72) | -1.09 (0.7) |
| Isthmus cingulate | -1.8 (0.67) | -0.24 (0.92) | -1.5 (0.7) |
| Precuneus | -2.14 (0.52) | -0.32 (0.91) | -1.99 (0.49) |
| Caudal anterior cingulate | -2.51 (0.4) | -2.78 (0.31) | 0.07 (1) |

**Supplementary Table S4:** Results of the unpaired group-wise t-tests of functional connectome node strength (averaged over the right and left homologues) between controls and TBI at 1 and 6 months and between TBI patients at 1 and 6 months.

**Supplementary Figures**

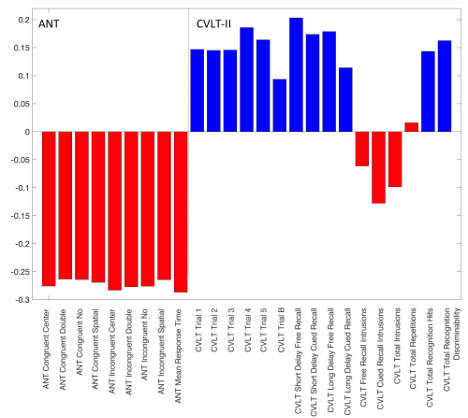

**Supplementary Figure S1:** The coefficients of the first component of the PCA analysis performed over all available neuropsychological data sets at all time points (N = 138). ANT scores are on the left and CVLT-II scores are on the right. Red indicates that sub-score’s values are smaller = better and blue indicates that sub-score’s values are higher = better.

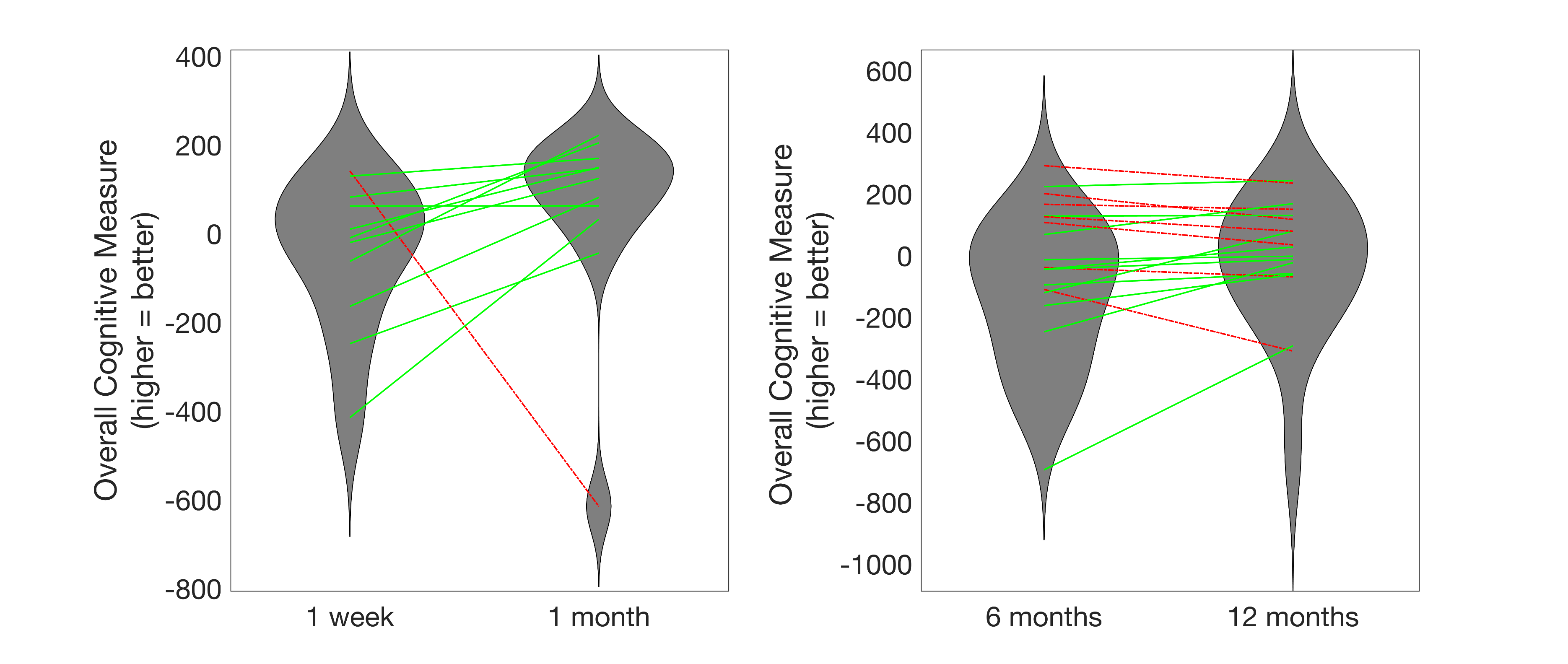

**Supplementary Figure S2:** Violin plots of the overall cognitive measure at each time point pair (Panel A: 1 week and 1 month and Panel B: 6 months and 1 year), along with the change in those scores over time. Lines represent individuals, where red indicates those whose overall cognition got worse over time and green indicates those patients whose overall cognition got better over time.

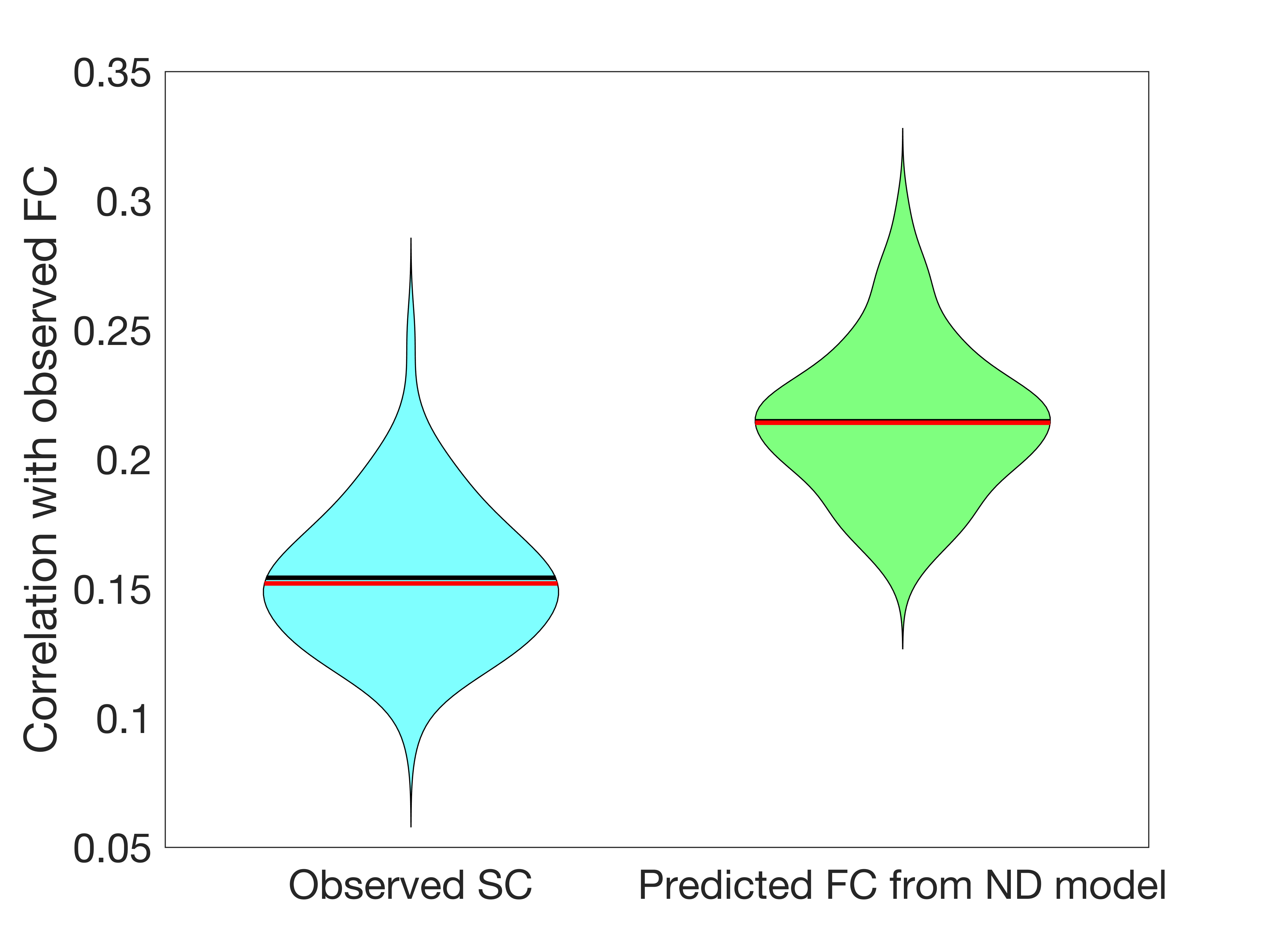

**Supplementary Figure S3:** Violin plots of the correlation between observed functional connectivity and observed structural connectivity (left, blue) and the Network Diffusion model’s predicted functional connectivity (right, green). The Network Diffusion model’s predictions had significantly higher correlations with observed functional connectivity than a model based only on structural connectivity (t = 42.3, p ≈ 0).

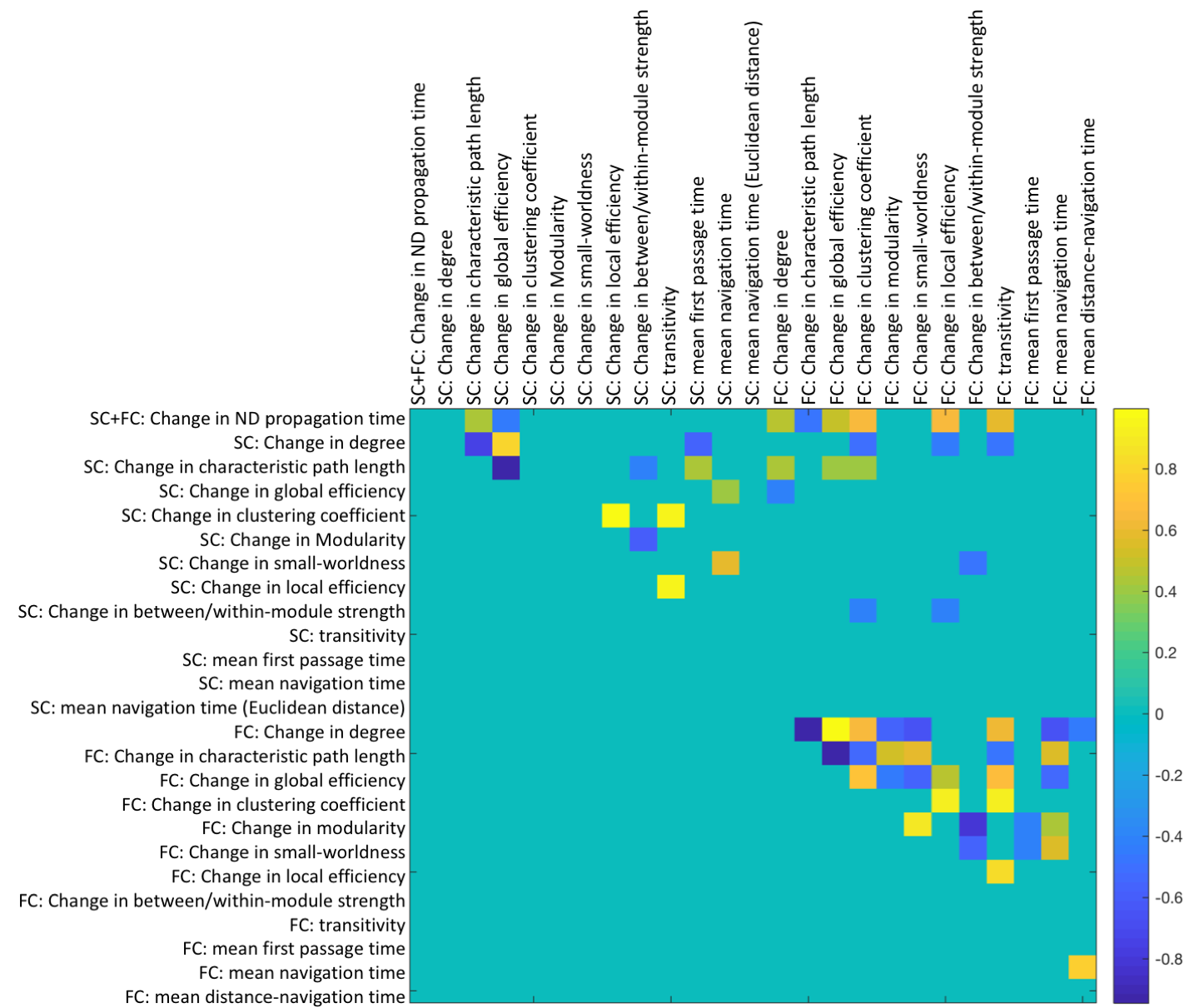
**Supplementary Figure S4:** Correlation matrix between changes ($\Delta m=m_{FU}-m_{BL})$ in FC and SC metrics between 1 and 6 months. Entries with non-zero elements were significant (uncorrected p<0.05).

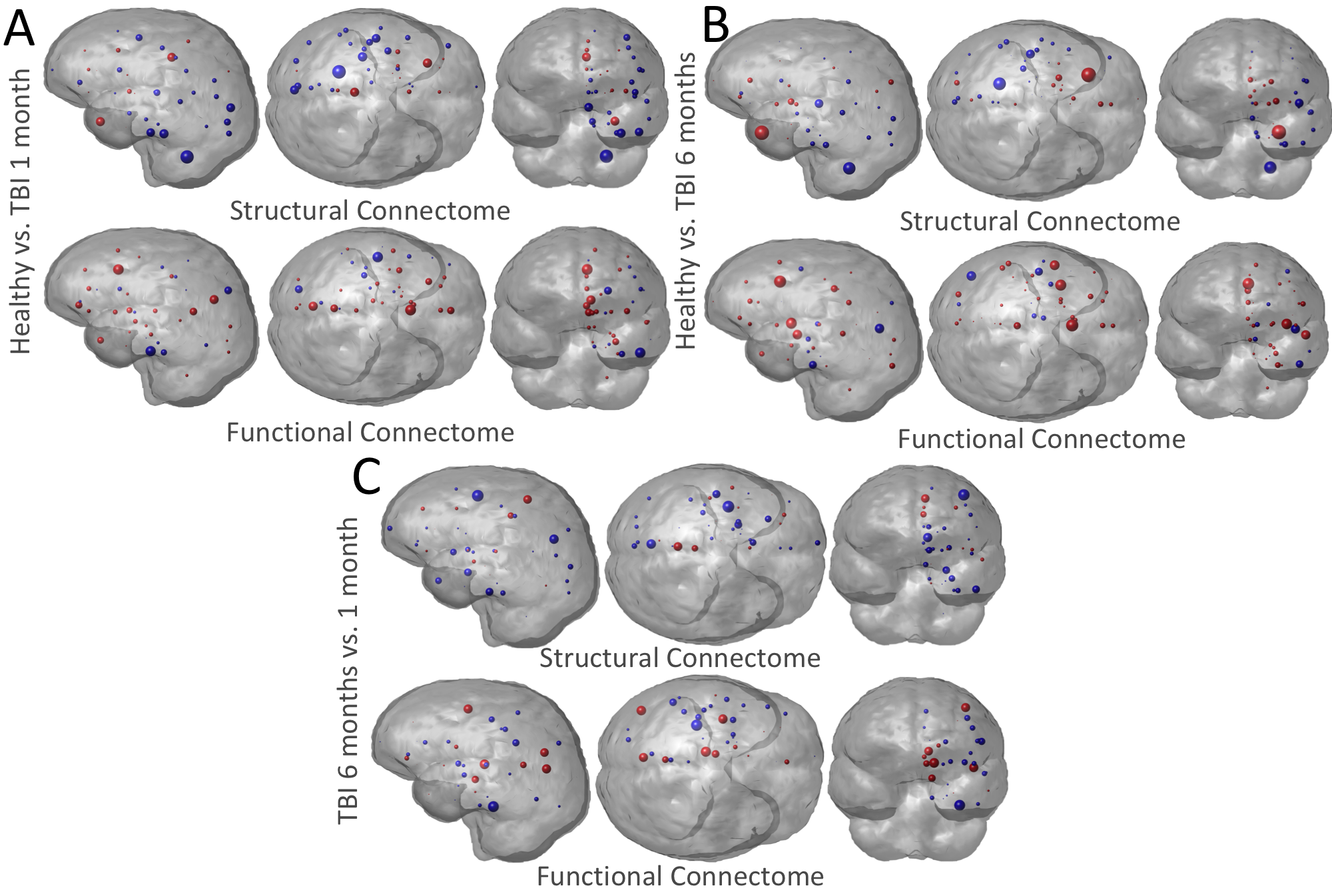

**Supplementary Figure S4:** Glassbrain visualizations of structural and functional connectome regional node strength differences (via group-wise t-statistics) between A) controls - TBI patients at 1 month b) controls - TBI patients at 6 months and C) TBI patients at 1 month - 6 months. Each region is represented by a sphere, where the diameter is proportional to the magnitude of the t-statistic and the color indicates the sign (red = negative, blue = positive). For example, in panel A the blue spheres indicate regions where the controls have higher node strength compared to the 1 month TBI patients. Results shown for left cerebral hemisphere only due to the combination of left/right region pairs.
